# Loser effects orchestrate dominance hierarchies in socially-controlled sex change

**DOI:** 10.64898/2026.03.16.712238

**Authors:** Haylee M. Quertermous, Chloé A. van der Burg, Kaj Kamstra, Simon Muncaster, Christine L. Jasoni, Culum Brown, Neil J. Gemmell

## Abstract

Socially-controlled sex changing fishes provide powerful model systems for investigating sexual development and phenotypic plasticity in both behavior and physiology. The remarkable sexual transformation these fishes undertake is strongly influenced by their position in dominance hierarchies. However, the behavioral mechanisms underlying hierarchical formation remain understudied, particularly among female groups. Here, we investigated the role of winner-loser effects among females in establishing social dominance in a female-to-male sex changing fish. Individuals with prior losing experiences were more likely to lose subsequent size-matched fights, demonstrating clear loser effects, while there was no evidence for winner effects. Initial mirror aggression and some prior fighting behaviors, particularly submission, significantly and positively correlated with aggression in size-matched fights and subsequent mirror aggression; however, contest outcomes were not altered by these factors. Additionally, mirror aggression increased significantly only in subjects that drew size-matched fights. These findings demonstrate complex fighting dynamics in female-female competition and confirm the presence of loser effects in a sequential hermaphroditic species. These effects may represent evolutionarily advantageous mechanisms underlying sex change, thereby offering further context for examining how social rank advantages drive sexual transition.

## Introduction

Sex change is predicted to be evolutionarily advantageous, as larger individuals that make the transition are expected to increase their reproductive success (i.e., size advantage hypothesis) (1, 2). However, social environment is also a well-established factor influencing sex change (3). Among the approximately 500 species of sequentially hermaphroditic fishes, it is typically socially dominant individuals that undergo sexual transition (4–6).

All three types of sex change (protogyny, protandry, and bidirectional) can be governed by social dynamics. Protogynous (female-to-male) sex change is frequently associated with polygynous mating systems, in which a single dominant male monopolizes a group of females (7). When the male is lost, the highest-ranking female rapidly escalates aggressive behavior to assume the male role and initiates sex change (4, 8). Similarly, in many protandrous and bidirectional species, transformation occurs in the most dominant individual within the group (3, 5, 6). In protogynous species, behavioral and physiological markers of sex change can emerge within minutes of male removal (4, 9), suggesting that dominant females may already be prepared to transition before social disruption occurs (10). Consequently, understanding female dominance hierarchies and the behavioral mechanisms that regulate them is essential for elucidating how sex change is primed and initiated.

Dominance hierarchies are a common social structure in both laboratory and wild populations (11, 12). Linear hierarchies prevail in many species, with a single top-ranked individual that dominates all others, a second that dominates all except the first, and so on (13, 14). Sequential hermaphroditic species often establish linear hierarchies, with social structure likely playing a critical role as individuals queue for the opportunity to change sex (9, 10, 15). These hierarchies can benefit groups by minimizing conflict, though the underlying behavioral processes driving their formation are still being unraveled (12–14). A key aspect of understanding social hierarchy formation involves examining how prior competitive interactions influence its development.

Winner-loser effects are common behavioral phenomena describing how prior fighting experiences influence hierarchy formation, particularly the likelihood of success in future encounters (16). Individuals that are more likely to lose following previous defeats display loser effects, whereas the opposite pattern characterizes winner effects. Winner-loser effects occur across many taxa but are not universal: some species exhibit both effects, only one, or neither (17). The duration of winner-loser effects also varies widely, ranging from less than 24 hours to more than a week in some cases. The ecological and evolutionary factors shaping the presence and strength of these effects remain an active area of research (18).

Winner-loser effects are thought to carry evolutionary significance by reducing conflict and promoting the stability of social hierarchies (14). Although these effects have been documented in various fish species (16), they have not yet been investigated in sequential hermaphrodites and have received limited attention in female hierarchies more broadly (18). Sex-changing species, therefore, present a unique opportunity to explore the cost-benefit dynamics associated with winner-loser effects. Success in contests may confer substantial fitness benefits, such as allowing sex change and improving access to mates and preferred territory, thereby increasing reproductive output (1). Conversely, contest engagement may incur high costs, including injury, heightened aggression from dominant individuals, or group eviction (19). Thus, winner-loser effects may function as important behavioral mechanisms that facilitate strategic decision-making during contest and ultimately shape social hierarchy formation in sequentially hermaphroditic fish.

In addition to the effects of previous interactions, individual traits are also hypothesized to play a significant role in the formation of social hierarchies (11, 20). Key attributes influencing group structure include age, sex, physical size, strength, and aggressiveness (20–22). When examining individual behavioral traits, it is also important to consider behavioral consistency, or personality. Many species exhibit stable individual differences in behavior over time (23), with aggressive behavior in particular being repeatable (22, 24). However, the extent to which social experience, individual traits, behavioral plasticity, and their interactions contribute to fight outcomes remains unclear.

Here, we investigate whether the protogynous New Zealand spotty wrasse (*Notolabrus celidotus*, hereafter “spotty”) displays winner-loser effects and the extent to which their aggressive behavior is plastic. Spotty are an emerging model system for studying protogynous sex change, with research spanning their ecology, behavior, and the physiological mechanisms underlying sex change (9, 25–30) (Figure 1). Our previous research has shown that spotty form linear hierarchies and that second-ranked individuals respond rapidly to the removal of the most dominant fish, quickly initiating aggressive “male-like” behavior and activating the brain’s social decision-making network (9), alongside molecular and endocrine changes in the gonad (29) (Figure 1). By focusing on female spotty, we investigate the behavioral mechanisms that facilitate hierarchy formation while also addressing the pervasive male bias in behavioral research (18), thereby contributing novel insights into the nature and extent of female-female interactions.

**Figure 1.**
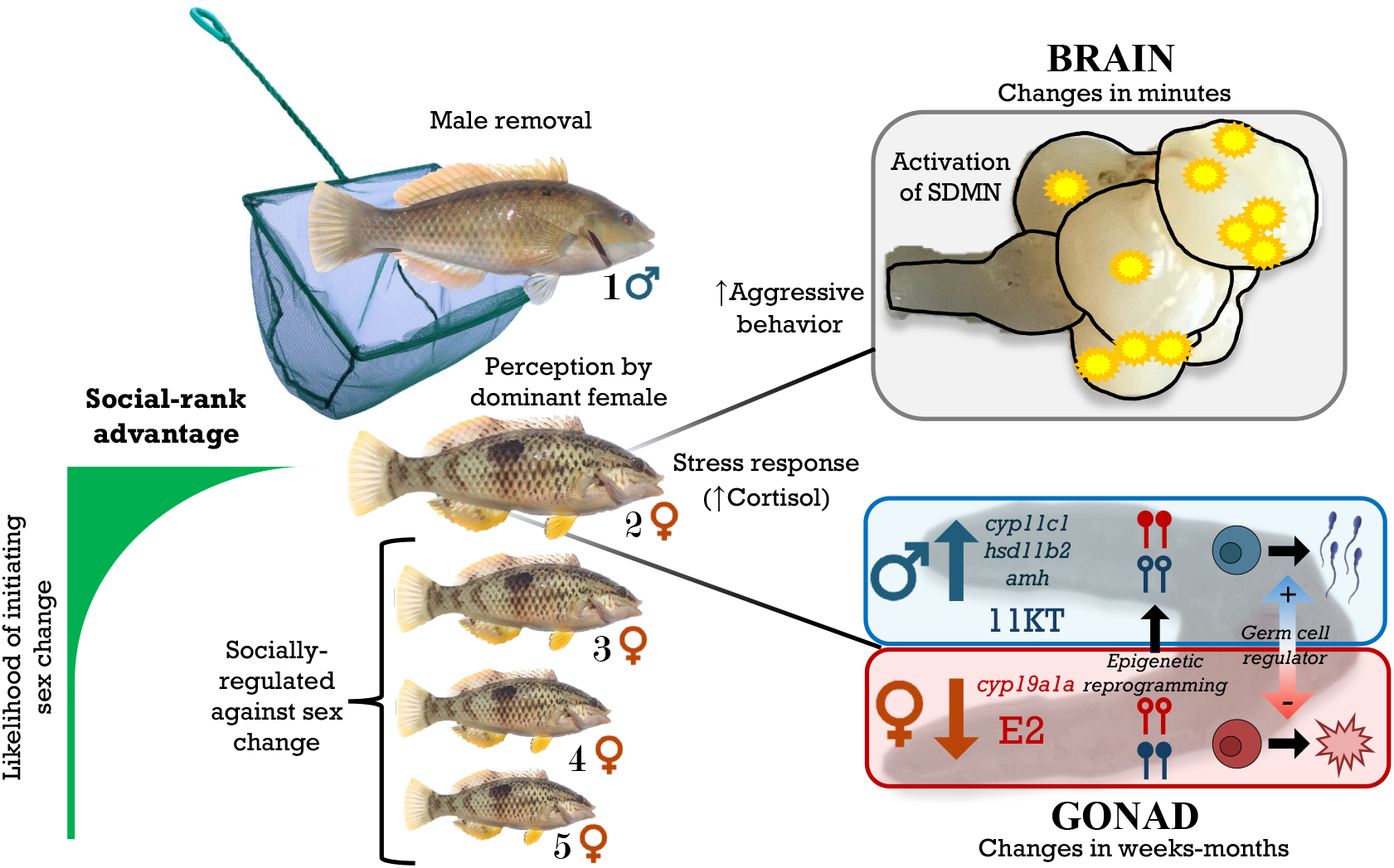
Initiation and process of sex change controlled by social rank. Perception of a social cue (e.g., absence of dominant male) triggers sex change in the dominant female, while lower-ranking females are socially inhibited from undergoing sex change. Behavioral changes rapidly occur in the dominant female with increased aggression and activation of the social decision-making network (SDMN) in the brain. Physiological responses include increased cortisol levels, which promotes the transition of ovary (red) to testis (blue) in the gonad. Molecular and hormonal shifts in the gonad involve (i) downregulation of aromatase (*cyp191a*) expression causing estrogen (E2) production and feminizing expression to decline, as well as ovarian atresia; (ii) upregulation of *amh* expression, a transcription factor (TF) and germ cell regulator that inhibits feminizing genes and promotes oocyte apoptosis while also promoting masculinizing and spermatogonial recruitment; (iii) upregulation of androgenic genes *cyp11c1* and *hsd11b2* to increase 11-Ketotestoterone (11-KT) production and support testicular development. Sexual fate and sex-specific expression is canalized via epigenetic reprogramming, changes in sexually dimorphic DNA methylation (shown as lollipop shape; open/unmethylated and filled/methylated). Modified from Todd, E.V., Ortega-Recalde, O., Liu, H., Lamm, M.S., Rutherford, K.M., Cross, H., Black, M.A., Kardailsky, O., Marshall Graves, J.A., Hore, T.A., Godwin, J.R., Gemmell, N.J., 2019. Stress, novel sex genes, and epigenetic reprogramming orchestrate socially controlled sex change. Science Advances 5, 1–15. https://doi.org/10.1126/sciadv.aaw7006.

To test for winner and loser effects, we employed a random assignment protocol (31): assigned winners consecutively interacted with three smaller opponents, whereas assigned losers interacted with three larger opponents (Figure 2A). Following these encounters, each focal fish was paired with a size-matched, socially inexperienced individual for a fourth and final contest (16). Fight outcomes were determined by quantifying submissive and aggressive behaviors displayed during interactions. We also assessed individual aggression using mirror tests (mirror aggression) conducted before and after the winner-loser manipulations (Figure 2B). This approach allowed us to evaluate whether baseline aggression influences fight outcomes, the extent to which aggression is a consistent (repeatable) individual trait, and whether aggression changes in response to social experience (behavioral plasticity).

**Figure 2.**
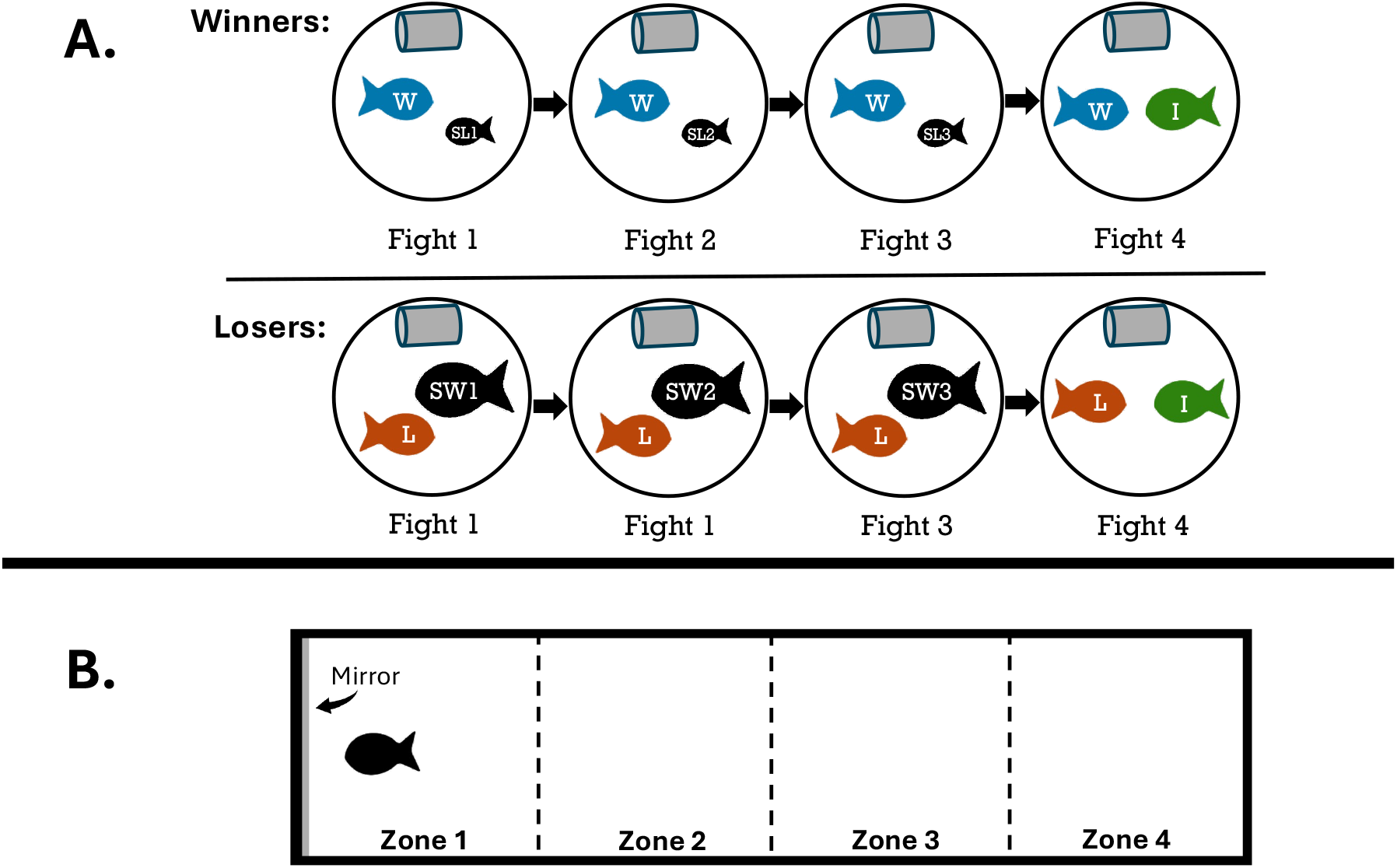
Experimental Design. (A) In the winner/loser experience phase, assigned winners engaged in three successive 20-minute contests against smaller standard losers (SL), while assigned losers fought three times against larger standard winners (SW). In the fourth contest, both groups were matched against a size-matched, inexperienced (I) fish to assess the effects of prior experience. (B) The individual aggression arena was divided into four evenly spaced zones, with a mirror placed at one end in Zone 1 to simulate a size-matched opponent.

We hypothesized that spotty would exhibit both loser and winner effects. Specifically, we predicted that individuals that lost initial fights would be more likely to lose the final fight, and individuals that won initial fights would be more likely to win the final fight. We further hypothesized that behavior during earlier fights would influence behavior in the final fight. Accordingly, we predicted that higher levels of submissive behavior during the initial fights would be associated with lower aggression in the final fight, while higher levels of aggressive behavior during the initial fights would be associated with higher aggression in the final fight. Finally, we predicted that mirror aggression would be positively associated with aggression in the final fight. We also predicted that mirror aggression would be plastic in response to fight outcomes, increasing in winners and decreasing in losers.

Social status is a primary factor in initiating sex change. However, a comprehensive hypothesis to guide further investigation of the behavioral mechanisms is yet to be established. Here, we propose the “social rank advantage hypothesis” to guide future research examining how individuals of varying social status are differentially primed, with higher status individuals having increase potential for sex change (Figure 1).

## Materials and Methods

### 1. Experimental Fish

Fish handling was performed in accordance with New Zealand National Animal Ethics Advisory Committee University of Waikato Protocol 1181. Spotty were captured using hook and line off the coast of Tauranga, Bay of Plenty, New Zealand (37.6878° S, 176.1651° E), between May 2023 and July 2024, and then held at the Coastal Marine Field Station, University of Waikato, Tauranga. Fish were housed in 400 L recirculating holding tanks to acclimate at least a couple of months prior to experimentation. Prior to experimental allocation, fish were anaesthetized through immersion in 600 ppm 2-phenoxyethanol; standard length (SL; ranging from 74-180 mm) and body weight (ranging from 5.4 – 120.3 g) were measured. Floy tags (Fine Anchor T-bar 5 cm, Hallprint T-Bar tags) were attached dorsally to enable individual identification. Fish were fed marine fish pellets (BioMar™) three times per week.

### 2. Experimental Design

#### a. Mirror Aggression

A standard mirror test was conducted to assessIndividual aggression in a 60 L glass tank (60 cm x 30 cm x 30 cm) (Figure 2B), filled with ∼30 L of water. Tanks were surrounded by opaque black tarp to block external visual stimuli. Self-adhesive flexible mirrors were fitted to the width of tanks and attached to one side. Fish were placed into the center of tanks, and behavior interactions with the mirror images recorded for 10 minutes using GoPro cameras (HERO Session or Hero 11 Black Mini models) mounted above tanks. None of the fish had previous mirror exposure. Mirror aggression was measured the day before (∼24 h) the four winner-loser effect fights and immediately after fights.

#### b. Winner-Loser Effects

Fish were randomly assigned one of three roles: loser, winner, or inexperienced. A random assignment protocol with size asymmetries was established to provide winning and losing experiences (31). Assigned losers underwent three consecutive 20-minute defeats against larger opponents (“standard winners”), followed by a fourth 20-minute size-matched fight with an inexperienced fish (Figure 2A). Similarly, assigned winners underwent three consecutive wins against smaller “standard losers” before the fourth size-matched fight. All opponents came from different holding tanks to ensure unfamiliarity.

Prior to fights, experimental fish were transferred from their home tanks into individual 20 L buckets. With all fish collected, each pair was introduced simultaneously into contest arenas, identical 60 L flexi tub buckets filled with ∼25 L of water. 100 mm wide DWV pipes, cut to 200 mm long and attached with 3M Command™ picture hanging strips to arena edges, acted as a shelter. GoPro cameras recorded interactions from above.

After each fight, fish were removed and returned to their 20 L bucket before being placed into a different arena for the next experience. Inexperienced fish were individually moved between arenas during the first three fights to control for handling effects. After the fourth fight, fish were immediately transferred to the mirror aggression tanks for behavioral assessment before being returned to their original tanks. All procedures were conducted between 09:00-15:00 NZST, with winner and loser fights alternating between morning and afternoon start times.

### 3. Behavior Analysis

All videos were analyzed using BORIS software (32). For mirror aggression trials, aggressive behaviors toward mirror images and location within four equally sized zones were analyzed over a 10 min period (Table S1). An aggression score was calculated by summing all aggressive behavior counts to the total seconds spent frontal swimming (i.e., one second equals one aggressive behavior).

For WL effects, aggressive and submissive behaviors were recorded, along with time spent in or behind the shelter (Table S1). Aggressive (mouth wrestling, chase, rush, attack, head bob, wiggle) and submissive behaviors (escape, flinch) were added to get an overall aggression and submission score. A dominance score was calculated using the formula: Dominance score = #Aggression / (#Aggression + #Submission) (similar to 9). A dominance score close to 1 indicates a highly dominant individual, while a score close to 0 is indicative of a highly submissive fish.

A fight was considered a win or loss if the winner displayed three or more aggressive behaviors or the loser showed three or more submissive behaviors relative to their opponent. If interactions were minimal or aggressive/submissive behaviors were evenly matched (<3 difference in the number of aggressive/submissive behaviors between opponents), the fight was recorded as a draw.

### 4. Statistical Analysis

All statistics were completed in RStudio Version 2025.05.0+496. Data frames were compiled from BORIS aggregated event files using the tidyverse meta-package.

Chi-squared tests were performed to assess differences in fight outcomes, with the expected probabilities set equally across three possible outcomes: win, lose, and draw. To account for small sample sizes and distributional assumptions, Monte Carlo simulations were used, running five independent chains with 1 million iterations (33).

General linear mixed models compared behavior measurements between fights, using lme4 and glmmTMB packages. Home tank of subject fish and fight arena were included as random effects. The best model distribution was determined using a custom function (compare_distributions). The function fits a series of candidate models with various distributional assumptions (Poisson, negative binomial, Gaussian, gamma, Tweedie, zero-inflated Poisson, and zero-inflated negative binomial) using glmmTMB, evaluated residuals using the DHARMa package, and ranked models based on (Akaike Information Criterion) AIC values (34, 35). Some behavior variables (duration in shelter, under shelter, in Zone 1, and in Zone 4) were square root transformed to better adhere to model assumptions.

Generalized linear mixed models, with subject’s home tank set as a random effect, investigated predictors of fight 4 aggression. Predictor variables included counts of submissive and aggressive behaviors across the previous three fights and mirror aggression scores before WL effects fights. mirror aggression after WL effects fights was also analyzed using submissive and aggressive behaviors from all four fights and mirror aggression before fights. Multicollinearity among predictors was assessed using the vif function from the car package. Model selection was achieved using backward stepwise selection. Following the identification of the best set of predictor variables, the best distribution for each model was determined using the compare_distributions function, as previously described.

Wilcoxon signed-rank tests evaluated differences between mirror aggression before and after WL effects trials, both for assigned winners and losers, and individuals categorized by win, lose, or draw of their fight 4 outcome.

## Results

### 1. Presence of Winner-Loser Effects

Two assigned losers drew all training fights and were removed from analyses (Table S2). The remaining assigned losers lost or drew the final size-matched fights (Figure 3A), significantly losing more and winning less than expected by chance (Table S3). Assigned winners experienced all outcomes (win, loss, draw) in size-matched fights (Figure 3B) and did not differ significantly from chance expectations (Table S3).

**Figure 3.**
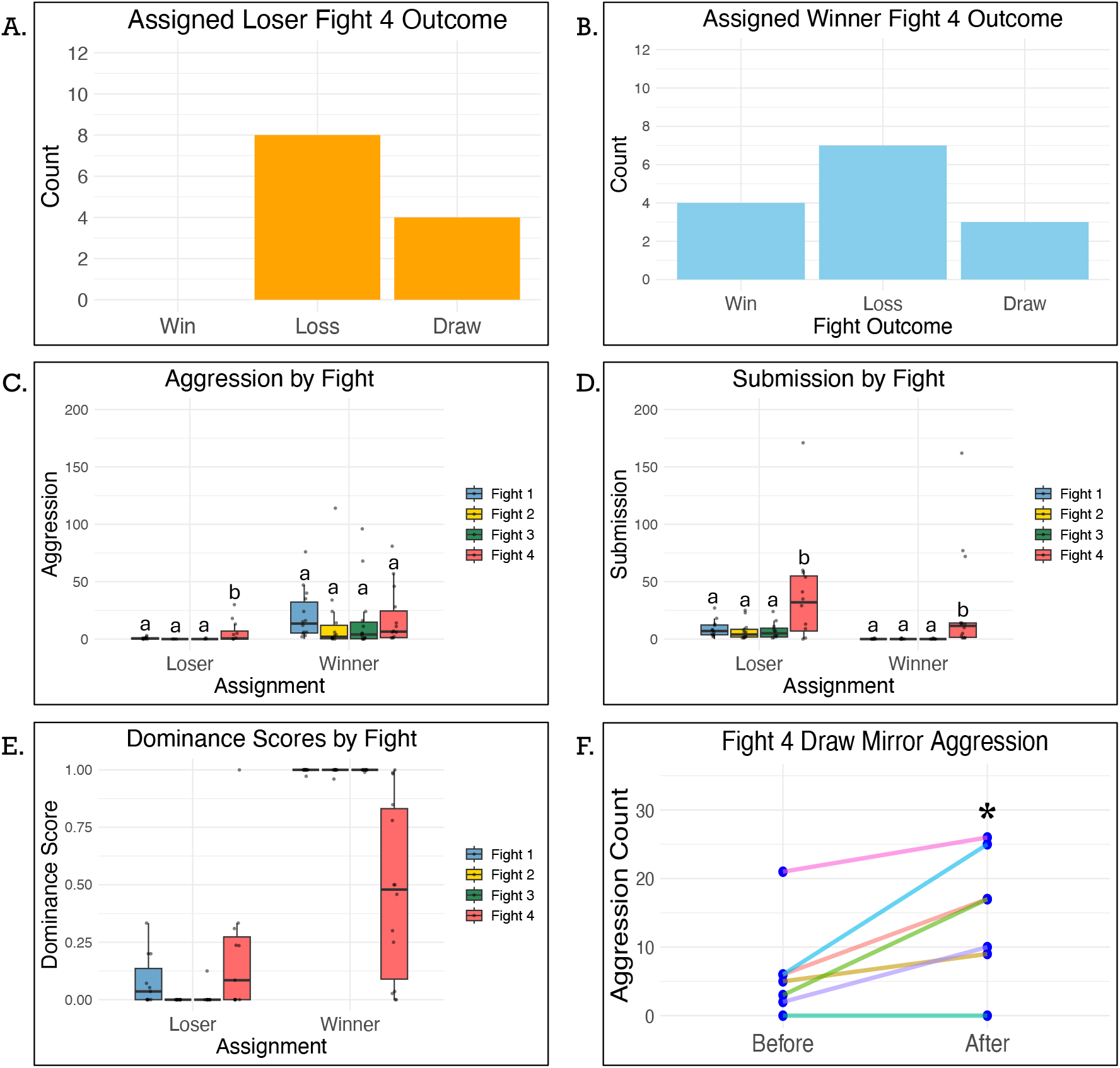
Fight 4 Outcomes and Fight Behaviors. (A) assigned losers lost or drew all fights against inexperienced size-matched individuals. (B) Assigned winners experienced all outcomes against inexperienced size-matched individuals in Fight 4. (C) Aggression count was significantly higher in Fight 4 for assigned losers (N = 12) but does not differ for assigned winners (N = 14). (D) Both assigned losers and winners had significantly higher submission in Fight 4. (E) Dominance scores were calculated with a formula including aggressive and submissive behaviors; they range between 0 and 1, with 0 indicating fully submissive behavior and 1 indicating fully aggressive behavior (F) Mirror aggression significantly increased after winner-loser effects trials for individuals that drew the final size-matched fight (N = 7) regardless of if they were assigned to be a winner or loser in the trials.

### 2. Mirror Aggression and Prior Fight Behavior Effects on Fight 4 Aggression

Assigned losers exhibited significantly higher aggression in fight 4 compared to the previous three fights, which did not differ significantly from one another (Figure 3C; Table S4). For Fight 2, the estimated coefficient and standard error from the model were extremely large. This is a consequence of there being no observed variation in aggression during this fight, as all subjects exhibited zero aggression. This lack of variation arises from the experimental manipulation, where all subjects were intended to lose and, consequently, did not display any aggression. Assigned winners showed no significant differences in aggression across all four fights (Figure 3C; Table S4).

Both assigned losers and winners showed significant increases in submission during fight 4 compared to fights 1-3 (Figure 3D; Table S4). Submission counts did not differ significantly among fights 1-3 for either group. Neither assigned winners nor losers had a significant difference in the time spent in or behind the shelter across the four fights (Table S5).

For assigned losers, fight 4 aggression was best predicted by mirror aggression before winner-loser fights and fight 1 submission (Table S6). Both fixed effects were positively and significantly associated with fight 4 aggression, with fight 1 submission showing a stronger effect than mirror aggression before trials. The random effect of tank accounted for minimal variance.

For assigned winners, the best predictors of fight 4 aggression were mirror aggression before trials, average fight 1-3 aggression, fight 1 submission, and fight 3 submission (Table S6). Among these, fight 3 submission had the largest and most significant negative effect, followed by Mirror aggression before trials also having a significant negative association. In contrast, average fight 1-3 aggression had a significant positive association with fight 4 aggression. Fight 1 submission showed a positive but non-significant effect. The random effect of tank explained a moderate amount of variance.

### 3. Mirror Aggression Repeatability

For assigned losers, mirror aggression after trials was best predicted by mirror aggression before trials and fight 1 submission (Table S6). Fight 1 submission had a significant positive effect on mirror aggression after trials, while mirror aggression before trials was positively associated but not statistically significant. The random effect of tank accounted for minimal variance.

For assigned winners, the best predictors of mirror aggression after trials were mirror aggression before trials, average fight 1-4 aggression, and fight 1, 2, and 4 submission (Table S6). All predictors had significant effects, with average fight 1-4 aggression having the strongest overall effect. Mirror aggression before trials and fight 1 submission were negatively associated with mirror aggression after trials, while the rest of the predictors had a positive effect. The random effect of tank explained little variance.

Mirror aggression before trials versus after trials did not differ significantly for assigned losers (V = 32.5, p = 0.638) or their opponents (V = 54, p = 0.576). Assigned winners also showed no significant change (V = 33, p = 1.000), although their previously inexperienced opponents exhibited a non-significant trend toward reduced aggression after trials (V = 62, p = 0.077). Mirror aggression did not differ significantly for individuals with a fight 4 lose outcome (V = 55.5, p = 0.506) or their winning opponents (V = 61.5, p = 0.593). Fight 4 winners (V = 3, p = 0.625) and their losing opponents (V = 9, p = 0.250) also showed no significant change. Interestingly, individuals with a fight 4 draw outcome showed a significant increase in mirror aggression (V = 0, p = 0.036) (Figure 3F), although their opponents did not differ (V = 19, p = 0.469).

## Discussion

Sequential hermaphrodites provide powerful model systems for investigating phenotypic plasticity at the behavioral and neurological level. Although social environment is pivotal in triggering sex change, the behavioral mechanisms influencing dominance hierarchies have been rarely explored in these species. Here, we demonstrate that loser effects exist in the protogynous spotty wrasse; however, winner effects were not detected. This work accentuates loser effects as a key social mechanism shaping dominance in sequential hermaphrodites, with important implications for sex changing opportunities, particularly in reinforcing social rank advantages in these fishes. While individual attributes can also influence responses to social interactions and rank determination (11, 20–22), they did not seem to play as large of an effect in this study. As predicted, individual aggression measured independently in a standard mirror test positively correlated with aggression performed in size-matched fights but did not alter overall outcomes. Surprisingly, and contrary to predictions, submissive behavior from a prior fight positively affected aggression in subsequent size-matched fights and post-fight mirror aggression levels. Equally unexpected, post-fight mirror aggression increased only in subjects that experienced a draw in the size-matched fight, regardless of whether due to minimal interaction or equal escalated fighting. Undetermined outcomes are often excluded from behavioral analyses, but manipulated unresolved conflicts in cichlids increase the likelihood of subsequent winning (36). These findings highlight the need to further investigate effects of ambiguous contest behavior. Although individual traits merit further exploration, our results primarily indicate loser effects exert the strongest influence on subsequent contest behavior.

Winner and loser effects are observed across a wide range of taxa, including humans (16, 18), although the magnitude of their influence remains debated (20). Early reviews suggested that loser effects are more prevalent and stronger than winner effects (16, 37), but a recent comprehensive meta-analysis found no consistent difference in magnitude across taxa (18), implying that both effects are equally important evolutionarily. Consequently, it is valuable to investigate the specific environmental factors favoring the presence or absence of both loser and winner effects (Figure 4).

**Figure 4.**
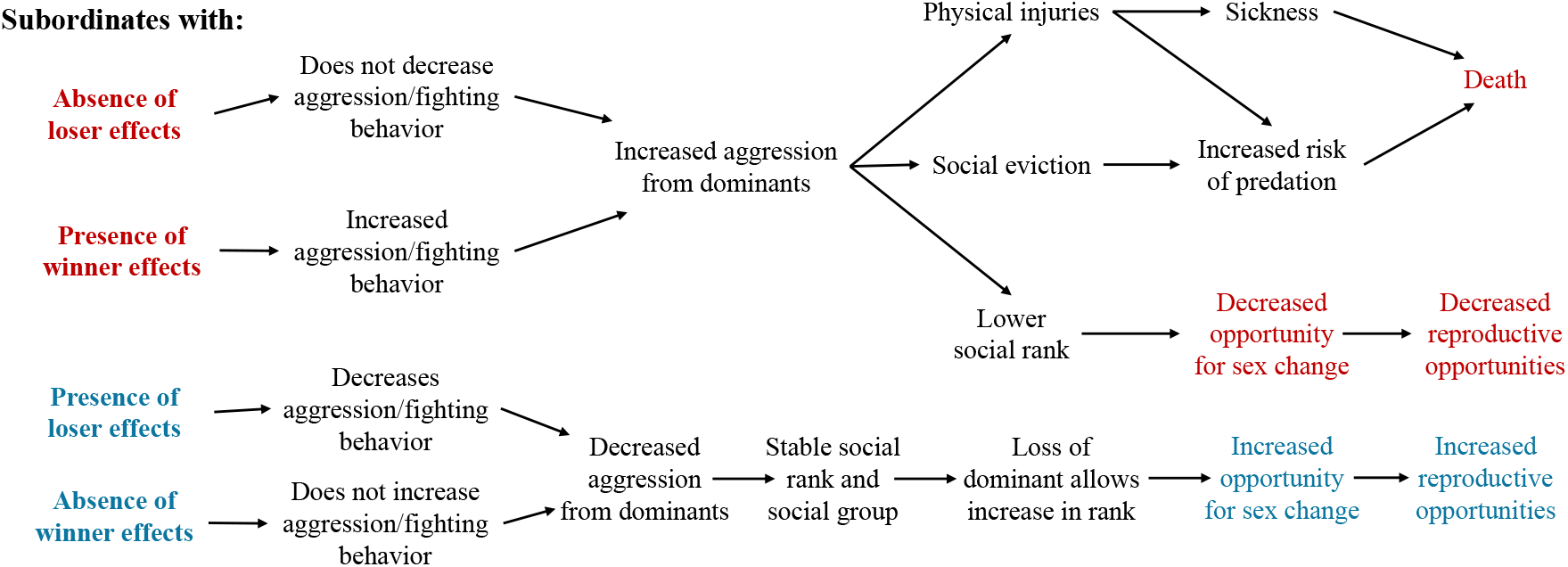
Costs and benefits of winner-loser effects. In protogynous sequential hermaphrodites that form linear dominance hierarchies, the absence of loser effects and presence of winner effects may be detrimental to subordinate individuals by failing to sufficiently regulate aggression, potentially leading to severe or fatal outcomes. Conversely, the presence of loser effects, as well as absence of winner effects, could reduce aggression and promote stable group formation, allowing subordinates to gradually ascend the hierarchy and increase their likelihood of undergoing sex change.

Loser effects are widely considered evolutionarily beneficial, especially when the costs of losing (e.g., injury, wasted energy, or lost time) are high (16). In protogynous fish, where a single dominant male suppresses sex change among a group of females: while male dominance is clear, interactions among females are also frequent and complex, as each individual is effectively queuing for the opportunity to change sex (9, 10, 38). In such systems, the absence of loser effects could be particularly detrimental with individuals motivated to maintain or improve their position in the hierarchy making them subject to increased aggression as part of social monitoring (19).

Here, our findings indicate loser effects serve a regulatory role stabilizing social hierarchies by modulating aggression. In linear dominance hierarchies, such social monitoring proves highly valuable. Absence of loser effects could have subordinates disregarding aggression mitigation and dominants escalating antagonism, thus heightening injury risk, lowering social rank, diverting energy from reproduction, and reducing sex change opportunity (Figure 4). Moreover, social ejection poses severe fitness costs, including elevated predation risk, challenges integrating into new social groups, and increased aggression from unfamiliar members (19, 39, 40). Our findings of loser effects in spotty emphasize the potential evolutionary stressors subordinates face within protogynous species.

However, winner effects, which typically emerge when winning provides substantial fitness gains, appear less beneficial in this context. In sequential hermaphrodites, this holds particular relevance as sex change confers considerable fitness benefits when individuals transition into dominant roles that increase their territory and reproductive output (1, 30, 41). Yet these effects may profit only those already near top ranks. For most group members, victories failing to secure dominant status likely yield minimal fitness returns.

Although social ranks were not determined in this study, future work should examine whether winner-loser effects vary across ranks. Many species, particularly sequential hermaphrodites, exhibit behavioral plasticity dependent on social rank (9, 42, 43); rank changes could similarly alter responses to contest outcomes. Both theoretical and empirical studies suggest age and experience modulate winner-loser effects (44, 45), though findings conflict necessitating further investigation. The strong loser effects observed in some studies (37) have also been suggested to reflect a bias toward younger subjects (44). Our study might not overcome this limitation; subjects were relatively small (90–139 mm SL, μ = 112.7 mm SL) compared to species’ maximum size (>230 mm SL) (26). While loser effects clearly shape spotty social behavior, exploring plasticity of these effects across age and rank, and their interactions, remains crucial for comprehending group social regulation (9, 19, 41).

Furthermore, understanding loser effects in protogynous species offer a powerful avenue for exploring how defeat shapes neural biology and inhibits sex change. As sex change mechanisms are increasingly unraveled, localized brain regions have emerged as key points of interest (46). The social decision-making network (SDMN), a highly conserved area associated with reproduction and aggression in vertebrates, shows rapid activation in spotty when introduced to permissive social environments (Figure 1) (9). Additionally, social defeat in male *Astatotilapia burtoni* elicits alternative coping responses correlated with activation of specific SDMN regions (47). In the protogynous *Thalassoma bifasciatum*, female social rank influences latency to male-like behaviors in permissive contexts, with dominants more readily initiating a change in behavior (10). Investigating how losing alters functioning of these brain regions, potentially “closing the gate” for sex change in subordinates, could illuminate pathways orchestrating sexual plasticity and further examine the social rank advantage hypothesis.

This study marks the first exploration of winner-loser effects in a sequential hermaphrodite, while also advancing our understanding of female contest behavior. A persistent literature bias favors male-male competition, especially in neural research on social rank, even though females routinely establish same-sex hierarchies and join mixed-sex ones (48, 49). Excluding females leaves a critical gap in dominance hierarchy studies, especially for revealing sex-specific dominance patterns within species.

## Conclusion

Overall, this study identifies loser effects as a fundamental social mechanism structuring dominance hierarchies in the New Zealand spotty wrasse (*Notolabrus celidotus*). Beyond shaping rank dynamics, these effects likely function as a key regulatory switch that constrains access to sex change, linking social experience directly to sexual fate in a protogynous vertebrate. By revealing how social feedback can gate one of the most extreme forms of phenotypic plasticity, this work reframes loser effects as an evolutionarily important control system rather than a by-product of competition. In doing so, it brings long-overlooked female hierarchies into focus and establishes sequential hermaphrodites as a powerful new model for uncovering how social behavior can drive developmental and evolutionary trajectories.

## Acknowledgments

Funding for this research was provided by a Marsden grant (UOO2115). The authors would like to acknowledge the assistance and technical support of Jana Longney, Bonnie Lewis, and Holly Ferguson and the statistical analyses assistance from Dr. Ryan Earley and Sebastián Alvarez Costes.

## Notes

**Competing Interest Statement:** The authors declare no conflict of interest.

https://doi.org/10.6084/m9.figshare.31350868

### Competing Interest Statement

The authors have declared no competing interest.

