## Supplementary Information for "Loser effects orchestrate dominance hierarchies in socially-controlled sex change"

**This PDF file includes:**

Tables S1 to S6
Legends for Movies S1 to S4
Legends for Datasets S1 to S6

**Other supporting materials for this manuscript include the following:**

Movies S1 to S4
Datasets S1 to S5

### Tables

**Table S1.** Ethogram of behaviors quantified in winner/loser effects and individual aggression trials.

| Experiment | Behavior | Behavior Type | Behavior Category | Description |
|---|---|---|---|---|
| **Individual Aggression** | Zone 1-4 | State | Location | Time spent in the four evenly split zones (30 x15cm) indicate how close fish were to the mirror (Zone 1 being the closest and 4 the farthest) |
| | Rush | Point | Aggression | Directed or accelerated movement towards the mirror (not resulting in a physical hit) |
| | Attack | Point | Aggression | Directed or accelerated movement towards the mirror resulting in a physical hit or a bite/nip. |
| | Wiggle | Point | Aggression | Vigorous wiggle of body while parallel to mirror (lateral display) |
| | Frontal Swimming | State | Aggression | Swimming along the mirror with contact from the snout (Balzarini et al., 2014) |
| **Winner/Loser Effects** | In Shelter | State | Location | Time spent in PVC pipe shelter |
| | Behind/Under Shelter | State | Location | Time spent between PVC pipe shelter and the arena wall or floor |
| | Mouthwrestling | Point/State | Mutual Aggression | An escalated form of aggression where opponents grip each other by the jaws and push and pull. |
| | Chase | Point/State | Aggression | Continued directed or accelerated movement towards an opponent while they are escaping |

| Experiment | Behavior | Behavior Type | Behavior Category | Description |
|---|---|---|---|---|
| | Rush | Point | Aggression | Directed or accelerated movement towards an opponent (not resulting in a physical hit) |
| | Attack | Point | Aggression | Directed or accelerated movement towards an opponent resulting in a physical hit or a bite/nip. |
| | Head bob | Point | Aggression | Rapid swinging of the head towards opponent |
| | Wiggle | Point | Aggression | Vigorous wiggle of body towards opponent (lateral display) |
| | Escape | Point | Submission | Change in swimming direction or acceleration away from opponent |
| | Flinch | Point | Submission | Small, sudden movement in response to aggression or presence of opponent |

**Table S2.** Aggregated fight outcomes. Possible outcomes include win (W), loss (L), and draw (D). Individuals P30 and R37 (red text) were removed from statistical analyses due to drawing all fights 1-3.

| Assignment | Fish ID | F1-F3 Outcomes | F4 Outcome |
|---|---|---|---|
| Losers | U07 | DDL | D |
| | Y33 | LDL | D |
| | R31 | LDL | D |
| | R28 | LDL | D |
| | P30 | DDD | L |
| | R37 | DDD | L |
| | R48 | DLD | L |
| | G44 | LDL | L |
| | P34 | LDL | L |
| | W12 | LLD | L |
| | R30 | LLD | L |
| | G41_9 | LLL | L |
| | P49 | LLL | L |
| | R32 | LLL | L |
| Winners | U19 | DWD | D |
| | R49 | WDD | D |
| | P29 | WDW | D |
| | P27 | DDW | L |
| | W49 | WDD | L |
| | P31 | WDD | L |
| | R38 | WDD | L |
| | R44 | WWD | L |
| | Y31 | WWW | L |
| | R39 | WWW | L |
| | P43 | WDW | W |
| | B22 | WWW | W |
| | R21 | WWW | W |
| | P48 | WWW | W |

**Table S3.** Chi-squared test results for Fight 4 outcomes.

| Assigned | Count | $X^2$ | p-value | Residuals |
|---|---|---|---|---|
| Loser | W – 0<br>L – 8<br>D – 4 | 8.000 | **0.017** | **W: -20000000**<br>**L: 2.0000000**<br>D: 0 |
| Winner | W –3<br>L – 7<br>D – 4 | 1.857 | 0.499 | W: -0. 7715167<br>L: 1.0801234<br>D: -0. 3086067 |

**Table S4.** General linear mixed model results of aggressive and submissive behavior across fights.

| Assigned | Behavior | Random Effects | Contrast | Estimate | SE | z.ratio | p.value |
|---|---|---|---|---|---|---|---|
| Losers | Aggression | Arena | Fight1 - Fight2 | 23.942 | 52688.317 | 0.000 | 1.000 |
| | | $S^2 = 9.81e\text{-}10$ | Fight1 - Fight3 | 2.197 | 1.209 | 1.817 | 0.265 |
| | | SD = 3.13 e-5 | **Fight1 - Fight4** | **-2.172** | **0.689** | **-3.152** | **0.009** |
| | | Tank | Fight2 - Fight3 | -21.744 | 52688.317 | 0.000 | 1.000 |
| | | $S^2 = 1.01e\text{-}9$ | **Fight2 - Fight4** | **-26.114** | **52688.317** | **0.000** | **1.000** |
| | | SD = 3.18e-5 | **Fight3 - Fight4** | **-4.369** | **1.168** | **-3.741** | **0.001** |
| | Submission | Arena | Fight1 - Fight2 | 0.235 | 0.474 | 0.495 | 0.960 |
| | | $S^2 = 1.91e\text{-}8$ | Fight1 - Fight3 | 0.282 | 0.474 | 0.594 | 0.934 |
| | | SD = 1.38e-4 | **Fight1 - Fight4** | **-1.532** | **0.464** | **-3.304** | **0.005** |
| | | Tank | Fight2 - Fight3 | 0.047 | 0.477 | 0.099 | 1.000 |
| | | $S^2 = 1.64e\text{-}9$ | **Fight2 - Fight4** | **-1.767** | **0.466** | **-3.789** | **<0.001** |
| | | SD = 4.04e-5 | **Fight3 - Fight4** | **-1.814** | **0.467** | **-3.885** | **<0.001** |
| Winners | Aggression | Arena | Fight1 - Fight2 | 0.676 | 0.587 | 1.151 | 0.658 |
| | | $S^2 = 5.34e\text{-}8$ | Fight1 - Fight3 | 0.308 | 0.604 | 0.510 | 0.957 |
| | | SD = 2.31e-4 | Fight1 - Fight4 | 0.285 | 0.581 | 0.490 | 0.961 |
| | | Tank | Fight2 - Fight3 | -0.368 | 0.602 | -0.611 | 0.929 |
| | | $S^2 = 0.478$ | Fight2 - Fight4 | -0.391 | 0.588 | -0.665 | 0.910 |
| | | SD = 0.692 | Fight3 - Fight4 | -0.023 | 0.580 | -0.040 | 1.000 |
| | Submission | Arena | Fight1 - Fight2 | -2.70e-07 | 1.52 | -1.78E-07 | 1.000 |
| | | $S^2 = 4.11e\text{-}9$ | Fight1 - Fight3 | -3.20e-07 | 1.52 | -2.11E-07 | 1.000 |
| | | SD = 6.41e-5 | **Fight1 - Fight4** | **-5.96** | **1.14** | **-5.21E+00** | **<0.001** |
| | | Tank | Fight2 - Fight3 | -5.00e-08 | 1.52 | -3.29E-08 | 1.000 |
| | | $S^2 = 2.52e\text{-}9$ | **Fight2 - Fight4** | **-5.96** | **1.14** | **-5.21E+00** | **<0.001** |
| | | SD = 5.02e-5 | **Fight3 - Fight4** | **-5.96** | **1.14** | **-5.21E+00** | **<0.001** |

**Table S5.** General linear mixed model results of time spent in and under the shelter across fights.

| Assigned | Behavior | Random Effects | Contrast | Estimate | SE | z.ratio | p.value |
|---|---|---|---|---|---|---|---|
| Losers | In Shelter | Arena<br>$S^2$ = 1.46e-9 SD = 3.82e-5<br>Tank<br>$S^2$ = 1.59<br>SD = 1.26 | Fight1 - Fight2 | -0.098 | 0.290 | -0.337 | 0.987 |
| | | | Fight1 - Fight3 | -0.313 | 0.286 | -1.096 | 0.692 |
| | | | Fight1 - Fight4 | 0.116 | 0.303 | 0.384 | 0.981 |
| | | | Fight2 - Fight3 | -0.215 | 0.281 | -0.767 | 0.869 |
| | | | Fight2 - Fight4 | 0.214 | 0.296 | 0.724 | 0.888 |
| | | | Fight3 - Fight4 | 0.430 | 0.287 | 1.494 | 0.441 |
| | SQRT Under Shelter | Arena<br>$S^2$ = 6.49e-9 SD = 8.05e-5<br>Tank<br>$S^2$ = 0.249<br>SD = 0.499 | Fight1 - Fight2 | 0.010 | 0.452 | 0.022 | 1.000 |
| | | | Fight1 - Fight3 | 0.132 | 0.465 | 0.285 | 0.992 |
| | | | Fight1 - Fight4 | -0.197 | 0.436 | -0.452 | 0.969 |
| | | | Fight2 - Fight3 | 0.122 | 0.466 | 0.263 | 0.994 |
| | | | Fight2 - Fight4 | -0.207 | 0.437 | -0.474 | 0.965 |
| | | | Fight3 - Fight4 | -0.330 | 0.448 | -0.736 | 0.882 |
| Winners | SQRT In Shelter | Arena<br>$S^2$ = 1.58e-9 SD = 3.97e-5<br>Tank<br>$S^2$ = 0.158<br>SD = 0.398 | Fight1 - Fight2 | -0.104 | 0.224 | -0.463 | 0.967 |
| | | | Fight1 - Fight3 | -0.264 | 0.218 | -1.210 | 0.621 |
| | | | Fight1 - Fight4 | 0.101 | 0.233 | 0.432 | 0.973 |
| | | | Fight2 - Fight3 | -0.160 | 0.213 | -0.749 | 0.877 |
| | | | Fight2 - Fight4 | 0.204 | 0.229 | 0.893 | 0.809 |
| | | | Fight3 - Fight4 | 0.364 | 0.223 | 1.637 | 0.358 |
| | SQRT Under Shelter | Arena<br>$S^2$ = 0.439<br>SD = 0.663<br>Tank<br>$S^2$ = 0.656<br>SD = 0.810 | Fight1 - Fight2 | -0.084 | 0.495 | -0.169 | 0.998 |
| | | | Fight1 - Fight3 | 0.493 | 0.558 | 0.884 | 0.813 |
| | | | Fight1 - Fight4 | 0.304 | 0.529 | 0.576 | 0.939 |
| | | | Fight2 - Fight3 | 0.577 | 0.552 | 1.044 | 0.724 |
| | | | Fight2 - Fight4 | 0.388 | 0.526 | 0.738 | 0.882 |
| | | | Fight3 - Fight4 | -0.189 | 0.590 | -0.320 | 0.989 |

**Table S6.** Results of general linear mixed models examining how behaviors from previous fights and individual aggression before winner/loser effects influence final aggression in Fight 4 and individual aggression after winner/loser effects.

| Assigned | Behavior | Tank Random Effect | Predictor | Estimate (β) | SE | z.ratio | p.value |
|---|---|---|---|---|---|---|---|
| Losers | Fight 4 Aggression | $S^2 = 1.28e-10$ SE = 1.13e-5 | **Intercept** | **1.015** | **0.263** | **3.865** | **<0.001** |
| | | | **IA Before** | **0.029** | **0.008** | **3.497** | **<0.001** |
| | | | **F1 Submission** | **0.054** | **0.012** | **4.302** | **<0.001** |
| | Individual Aggression After WL Effects | $S^2 = 8.24e-10$ SE = 2.87e-5 | **Intercept** | **2.224** | **0.319** | **6.964** | **<0.001** |
| | | | **IA Before** | 0.006 | 0.006 | 0.905 | 0.366 |
| | | | **F1 Submission** | **0.0589** | **0.020** | **2.873** | **0.004** |
| Winners | Fight 4 Aggression | $S^2 = 1.94$ SE = 1.39 | **Intercept** | **1.327** | **0.627** | **2.117** | **0.034** |
| | | | **IA Before** | **-0.024** | **0.012** | **-2.044** | **0.041** |
| | | | **Avg F1-F3 Aggression** | **0.047** | **0.021** | **2.231** | **0.026** |
| | | | F1 Submission | 2.336 | 1.478 | 1.580 | 0.114 |
| | | | **F3 Submission** | **-3.591** | **0.929** | **-3.866** | **<0.001** |
| | Individual Aggression After WL Effects | $S^2 = 4.07e-10$ SE = 2.02e-5 | **Intercept** | **2.056** | **0.182** | **11.302** | **<0.001** |
| | | | **IA Before** | **-0.036** | **0.009** | **-4.061** | **<0.001** |
| | | | **Avg F1-F4 Aggression** | **0.035** | **0.005** | **7.368** | **<0.001** |
| | | | **F1 Submission** | **-1.577** | **0.481** | **-3.279** | **0.001** |
| | | | **F2 Submission** | **1.002** | **0.237** | **4.228** | **<0.001** |
| | | | **F4 Submission** | **0.009** | **0.002** | **4.264** | **<0.001** |

**Movie S1.** Winner-loser effects contest clip of the winner fish defending the shelter with aggressive behaviors (rushes, attacks) and the loser fish displaying submissive behaviors (escapes).

**Movie S2.** Winner-loser effects contest clip of the winner fish behaving aggressively toward (rushing, attacking) the loser fish while it submits (escapes).

**Movie S3.** Winner-loser effects contest clip of an escalated fight with opponents reciprocating aggressive behaviors (attacks, mouthwrestling).

**Movie S4.** Individual mirror aggression trial of the subject fish displaying aggressive behaviors (attacks, frontal swimming) toward its mirror image.

**Dataset S1.** BORIS-aggregated behavioral events from all winner–loser effect contests. Data include observation ID, observation date, video length, subject identifiers, and all recorded behaviors, including behavior category, behavior type, start time, stop time, and duration (when applicable).

**Dataset S2.** Winner–loser contest information. Data include observation ID, whether winner or loser effects were tested, fight number (1–4), contest arena, Fish IDs of assigned contestants (assigned winner, assigned loser, and inexperienced fish), fight outcome and escalation as determined by the researcher, fight winner and loser (linked to BORIS subject identifiers and Fish ID), and additional notes.

**Dataset S3.** BORIS-aggregated behavioral events from all individual mirror aggression trials. Data include observation ID, observation date, subject identifiers, and all recorded behaviors, including behavior category, behavior type, start time, stop time, and duration (when applicable).

**Dataset S4.** Individual mirror aggression trial information. Data include observation ID, time of measurement (before or after the winner-loser effects session), mirror arena, and Fish ID.

**Dataset S5.** Individual fish information. Data include experimental fish metrics (phenotype, weight, and length), original tank, and additional notes.
